# Dual representation of lexical uncertainty in human cortex

**DOI:** 10.1101/2025.08.08.669343

**Authors:** P. Guilleminot, B. Morillon

## Abstract

Prediction under uncertainty is a hallmark of human cognition(Friston, 2010), yet how the brain represents uncertainty remains unresolved(Bach and Dolan, 2012). Speech provides a powerful test case: lexical predictions involve thousands of alternatives, making classical Shannon entropy an ambiguous measure of uncertainty. Here we explored a generalization of Shannon entropy, the Rényi entropy family (Rényi, 1961), combined with information-theoretic tools and whole-brain intracranial EEG during naturalistic listening. Word-related neural activity was best explained by two complementary forms of uncertainty: dispersion, reflecting the number of plausible alternatives, and strength, reflecting the probability of the most likely word. These uncertainties were consistently encoded in each participant by distinct neural populations, with opposing computational goals. Dispersion shapes anticipatory sensory activity and dampens responses to unexpected inputs, consistent with Bayesian optimization(Pouget et al., 2013). Strength instead emerges only for surprising words in lexical circuits, amplifying prediction errors and promoting information seeking. Despite bidirectional exchange between these two populations, their dynamics are dominated by local computations(Heilbron et al., 2022), challenging purely hierarchical models of predictive processing. Our findings reveal that the brain approximates large lexical probability distributions using multiple summary statistics, providing a flexible, low-dimensional representation of uncertainty in complex cognitive tasks and advancing principles of predictive processing across cognition.

## Introduction

To adapt to a dynamic environment, the brain relies on predictions generated by a continuously updating model of the world (Knill and Pouget, 2004; Heilbron and Chait, 2018). During speech perception, these predictive processes are crucial to anticipate upcoming words throughout an ever changing context (Kalikow et al., 1977; Lau et al., 2008). Recent advances in large language models (LLMs) offer a promising avenue for studying this process. Trained on vast corpora, LLMs generate context-sensitive probability distributions over upcoming words, aligned with neural signals (Schrimpf et al., 2021; Caucheteux et al., 2023). As such, LLMs can provide computational observer models for speech, as commonly used in other statistical sequence learning tasks (Meyniel, 2020; Badre et al., 2012; Stern et al., 2010). Two key computational quantities derived from predictive probability distributions are: (1) uncertainty, the amount of unknown information about the next input (Shannon, 1951), and (2) surprisal, the amount of information actually conveyed by that input (Shannon, 1948). Both are computed in the human brain and are thought to have complementary roles during cognitive processing: (1) uncertainty represents the confidence in the prediction and weights the incoming inputs, while (2) surprisal represents the prediction error and drives the updates of the internal model (Stern et al., 2010; Frank et al., 2015; Meyniel et al., 2015; Heilbron et al., 2022; Donhauser and Baillet, 2020; Song et al., 2024).

Traditionally, uncertainty over discrete probability distributions, such as lexicon-bounded predictions, is quantified using Shannon entropy, a foundational measure of unpredictability grounded in information theory (Shannon, 1948) and widely adopted in neuroscience (Donhauser and Baillet, 2020; Friston, 2010; Bach and Dolan, 2012; Garner, 1962; Heilbron et al., 2022). However, when reduced to a single summary statistics, the concept of uncertainty can be ambiguous. For instance, when watching a weather forecast, uncertainty may arise from the number of plausible (i.e. non-null probability) possibilities (rainy: 12%, sunny: 12%, cloudy: 76%) or from the lack of a clear dominant prediction (rainy: 50%, sunny: 50%, cloudy: 0%), with both situations described by the same Shannon entropy value (*H* = 1.0) (Fig.6A,B,D,E). While Shannon entropy can already be ambiguous with as few as three alternatives, it is particularly ill-suited to lexical uncertainty, where thousands of alternatives may exist. To address this issue, we used a generalization of Shannon entropy: the Rényi entropy family (Rényi, 1961). It introduces a parameter, the Rényi order *α*, that modulates the weight given to unlikely events, thereby offering several measures of uncertainty. Although Rényi entropies have been applied in fields such as computer science and signal processing (Baraniuk et al., 2002; Sahoo and Arora, 2004), they have yet to be used to characterize the neural representation of uncertainty during naturalistic cognition. Combined with information-theoretic tools and human intracranial recordings, we examined which forms of uncertainty are represented by the brain during naturalistic speech perception, and what role they play in processing words.

Our results reveal that two complementary forms of uncertainty support speech processing, represented in each participant by two separate neural populations: the dispersion across plausible alternatives, which estimates the number of plausible upcoming words (i.e. words with a non-null probability) regardless of their actual probability value; and the strength of the dominant prediction, i.e. the probability of the most likely upcoming word. Rather than encoding Shannon entropy or the full Rényi family, the brain processes these two complementary summary statistics to approximate the large lexical probability distribution. This offers a general and flexible solution to the representation of discrete probability distributions during a complex cognitive task. Critically, dispersion dominates at the sensory level, while strength intervenes in higher lexical processing, indicating that distinct forms of uncertainty selectively contribute to different computations within the speech processing hierarchy. Notably, both forms of uncertainty interact synergistically with surprisal at similar latency, but in opposite direction: dispersion dampens the responses to unexpected inputs at early sensory stages, whereas strength amplifies them at later lexical stages. We further demonstrate the existence of bidirectional information flow between these two functional clusters. Predictive processing in language thus emerges not from a single hierarchical chain, nor from a structure built on separated predictive circuits, but from a distributed architecture where complementary local computations shape predictions and error propagation refines them.

## Results

### The brain does not represent Shannon entropy

To characterize how the human brain encodes lexical uncertainty during speech comprehension, we recorded intracranial EEG from 33 epileptic patients (5126 channels) as they listened to an audiobook (Fig.1A) and epoched data to word onsets (N=1438). For each word, we generated probability distributions using an LLM (Martin et al., 2019) and computed Rényi entropies (Rényi, 1961), spanning *α* values from 0.01 to 10 (N = 20; Fig.1A). We used Gaussian Copula Mutual Information (GCMI) to relate neural data to entropy values (Fig.1A)) (Ince et al., 2017). Around 9% of channels (456/5126) showed statistically significant Mutual Information (MI) with at least one type of Rényi entropy (1000 permutations, *p* = 0.05, cluster-corrected over time). Across significant channels, MI time-courses showed three peaks around word onset (−50, 100, 300 ms) with sustained activity up to 1 s (Fig.1B). Both peaks and sustained activity were modulated by the Rényi order *α*. Critically, Shannon entropy (*α* = 1) was not the strongest measure. Instead, on average across significant channels, lower-*α* entropies consistently yielded higher MI with neural data, a pattern robust across participants (31/33; Fig.9B). Neural activity thus preferentially tracks the dispersion of plausible outcomes.

**Fig. 1.**
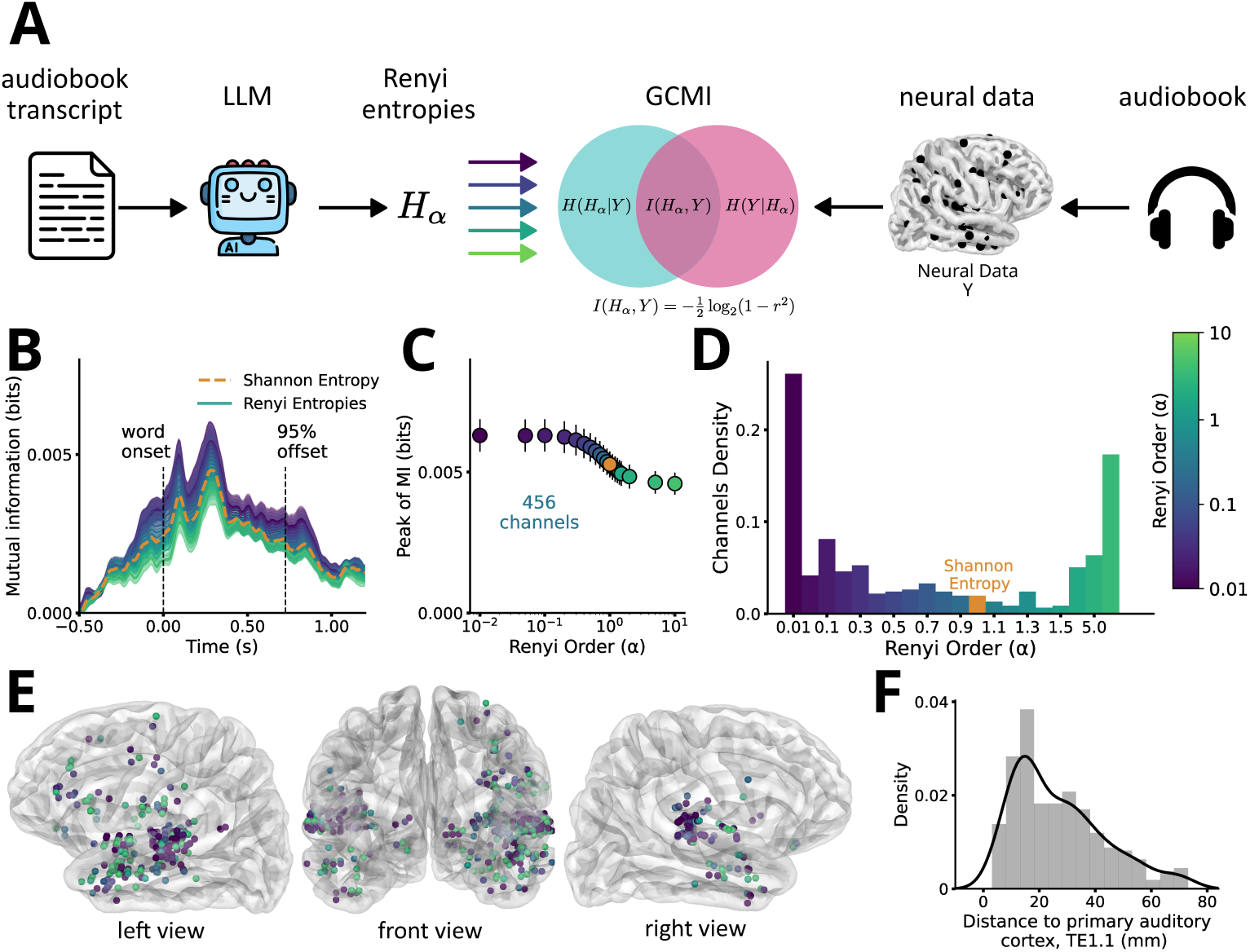
Neural representation of lexical uncertainty. **(A)** Analysis pipeline. Left: Transcript of an audiobook processed through an LLM to extract word probability distributions at each word onset. 20 Rényi entropies (*H_α_*) are computed for each distribution. Right: Intracranial EEG recorded during audiobook listening, epoched to word onset. Middle: Mutual information (MI) between Rényi entropies and epoched neural data, computed across words using Gaussian copula (GC) at each timestep between *−*500 ms and 1200 ms. **(B)** Time course of MI averaged across 456 significant channels, for the different Rényi orders (*α*, color gradient). **(C)** MI peak as a function of Rényi order, averaged across channels. **(D)** Distribution across channels of the Rényi order yielding maximum MI. **(E)** Cortical distribution of channels as a function of the Rényi order yielding maximal MI. **(F)** Histogram of distances from each channel to primary auditory cortex (posteromedial Heschl’s gyrus, TE1.1).

At the single-channel level, a more complex picture emerged: the *α* yielding the maximal MI peak per channel did not cluster around a single value, but instead fol-lowed a bimodal distribution with modes at low and high *α* (Fig.1D). A searchlight analysis showed that these different uncertainty coding schemes were distributed across cortex, with a concentration near auditory regions (Fig.1E-F). Thus, the brain does not represent Shannon entropy, but instead encodes lexical uncertainty heterogeneously across neural sites.

### Distinct neural populations encode distinct uncertainties

To characterize the heterogeneity in preferred uncertainty measures across neural sites, we applied a soft clustering method on MI peaks across *α* values. This revealed a simple two-cluster structure, each corresponding to a specific form of uncertainty (Fig.2). One cluster (248 channels) exhibited maximal sensitivity to low-*α* Rényi values, i.e. dispersion, the uncertainty relative to the number of plausible words (Fig.2A-C). The other (164 channels) was tuned to high *α* values, i.e. strength, the probability of the most likely word. Both clusters were present in all participants (Fig.2D-F).

**Fig. 2.**
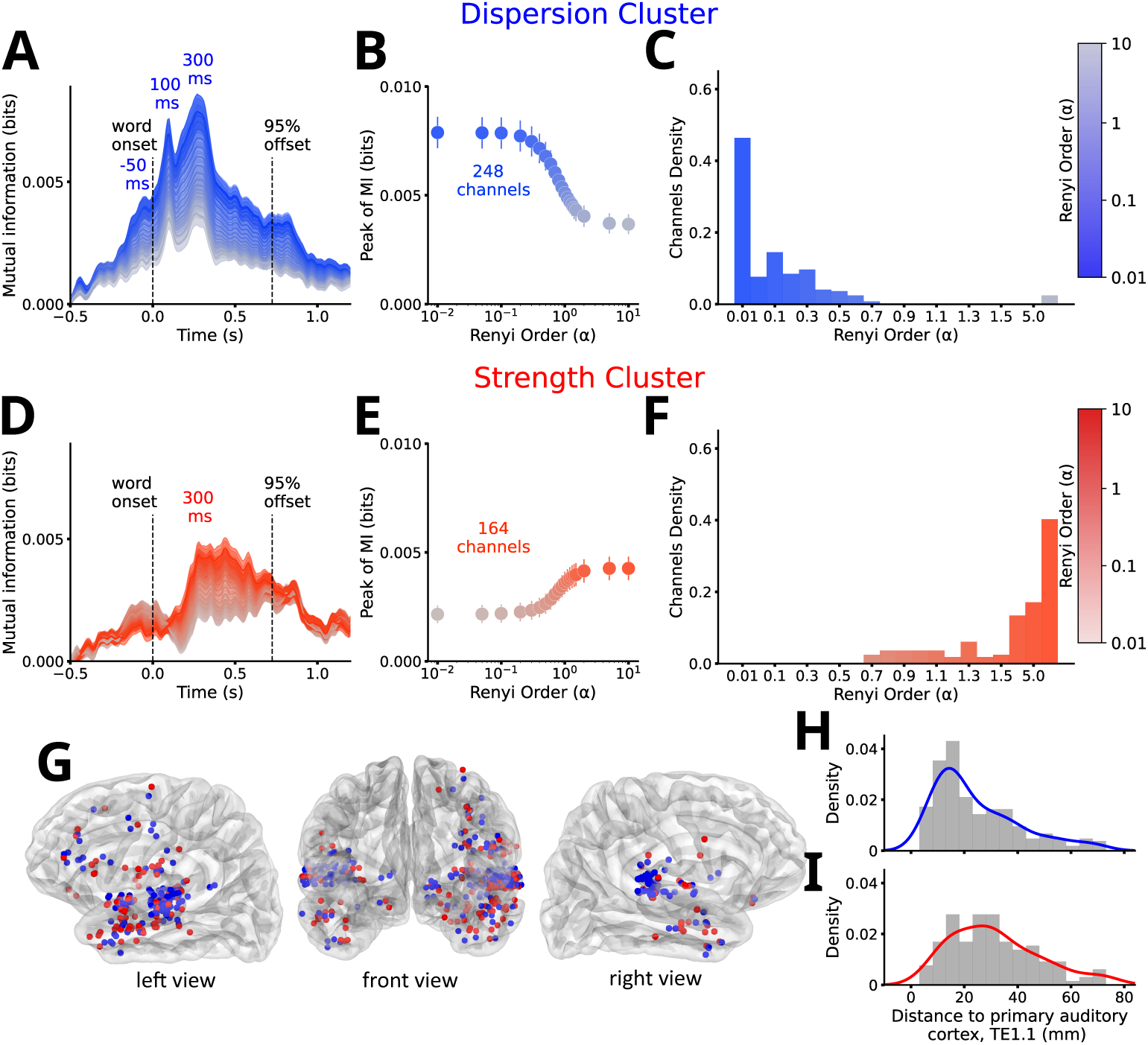
Dispersion and Strength clusters of channels. **(A-C)** Dispersion cluster (*n* = 248 channels). **(A)** Grand average of MI time courses across channels for different Rényi orders (*α*, color gradient). **(B)** MI peak as a function of Rényi order, averaged across channels. **(C)** Histogram of Rényi order yielding maximum MI with neural data, across channels. **(D-F)** Strength cluster (*n* = 164 channels). Same conventions as A-C. **(G)** Cortical distribution of the Dispersion (blue) and Strength (Red) clusters. **(H,I)** Histogram of distance to primary auditory cortex (posteromedial Heschl’s gyrus, TE1.1) across channels in the (**H**) Dispersion and (**I**) Strength clusters.

The dispersion cluster displayed all three peaks of the overall MI time-course together with sustained activity (Fig1B). In contrast, the strength cluster was dominated by the later 300-ms peak, which remained elevated before gradually decaying up to 1s (Fig1D). Given their similarity with known auditory responses (Sanders et al., 2002), these temporal profiles suggest that the dispersion cluster processes early sensory information, whereas the strength cluster supports higher-level lexical processing.

Anatomical analysis confirmed this division: the dispersion cluster was mostly localized near peri-auditory lower-level sensory areas, while the strength cluster was predominantly found in associative cortical regions (Fig.2G). Their distance from pri-mary auditory cortex differed significantly (two-tailed t-test, *t* = −4.01, *df* = 340, *p <* 10^−4^; Fig.2H,I), indicating that they occupy distinct stages of the speech processing hierarchy.

### Dispersion and strength interact with surprisal

We next disentangled the functional contributions of dispersion and strength clusters using Partial Information Decomposition (PID) (Williams and Beer, 2010) (Fig.3A,C-F). In both clusters, neural activity was partly redundantly explained by the two measures, consistent with their high mutual information (Fig.7A-C). Each cluster also carried unique information about its preferred uncertainty measure, except for a small onset-related contribution of dispersion in the strength cluster (t-tests: *p <* 0.05 FDR-corrected for multiple comparisons). Importantly, neither cluster showed synergy between dispersion and strength, indicating that the two statistics do not interact in either cluster. Overall, these results reveal a well-separated encoding of strength and dispersion within distinct neural populations (Fig.3F).

**Fig. 3.**
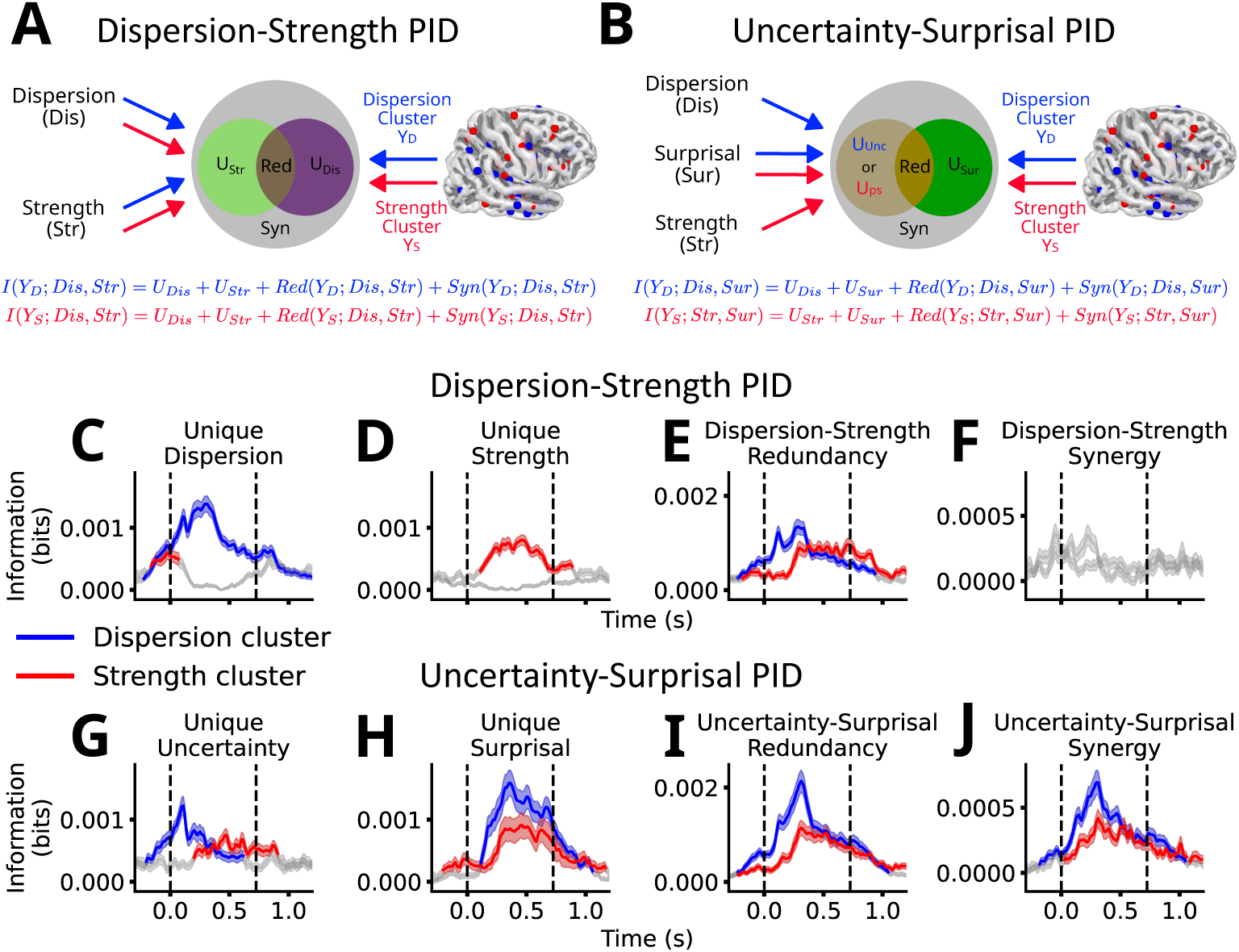
Partial Information Decomposition (PID) of the joint mutual information between neural data and LLM features. **(A)** Dispersion-Strength PID analysis framework: For each channel cluster (dispersion, blue; strength, red), the joint mutual information between neural data, dispersion (Rényi entropy with *α* = 0.01) and strength (Rényi entropy with *α* = 10) is com-puted. This is decomposed into the information about the neural data carried uniquely by dispersion, uniquely by strength, redundantly by both, and synergistically when combined. **(B)** Uncertainty-Surprisal PID analysis framework: For each channel cluster, the joint mutual information between neural data, surprisal (the unexpectedness of the observed word given the context) and the pre-ferred uncertainty (*α* = 0.01 or *α* = 10) for that cluster is computed. **(C,D,E,F)** Timecourses of the Dispersion-Strength PID components. **(G,H,I,J)** Timecourses of the Uncertainty-Surprisal PID components. Bold lines represent statistical significance compared to baseline (ttests: *p <* 0.05, FDR-corrected for multiple comparisons).

We further asked whether the brain also tracks the probability of the second most likely word. This measure carried no unique information in either cluster (p¿0.05 FDR-corrected), showing that neural activity does not represent the uncertainty carried by each candidate beyond the most likely. Rather, uncertainty is summarized only via dispersion and strength (Fig.10A,B,E-L).

Because the interaction between uncertainty and surprisal is central to predictive processing, we applied the PID between uncertainty and surprisal (Fig.3B, G-J). In their respective clusters, dispersion contributed uniquely from before word onset onward, whereas strength emerged only later, from ∼ 200 ms (Fig.3G). Surprisal was present in both clusters but emerged earlier in the dispersion cluster (∼ 100 ms) than in the strength cluster (∼ 200 ms), with both peaking around 300 ms (Fig.3H). In addition to redundancy, uncertainty and surprisal also exhibited synergy in both clusters, with clear peaks at ∼ 300 ms, demonstrating that dispersion and strength synergistically interact with surprisal at the same latency, despite their anatomical and functional dissociation (Fig.3I,J).

### Dispersion dampens, strength amplifies surprisal

The PID analysis revealed strong synergy between uncertainty and surprisal in both clusters, but did not characterize the direction of interaction. To test this, we orthogonalized uncertainty and surprisal across words, yielding four categories (quadrants; Fig.4A,C) and analyzed the event-related potential (ERP) for each.

**Fig. 4.**
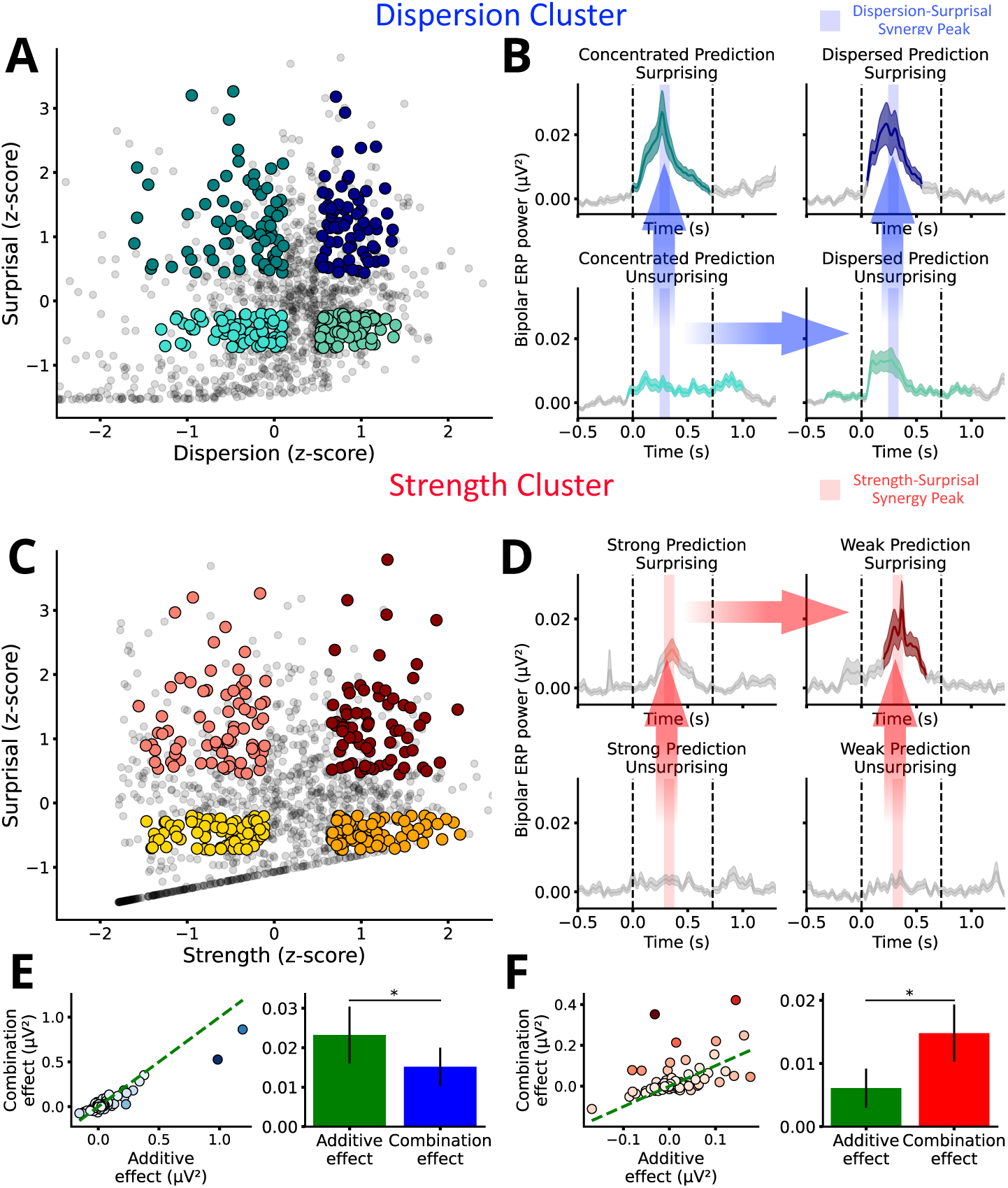
Evoked responses in the the Dispersion and Strength channel clusters as a func-tion of surprisal and uncertainty. **(A-B)** Dispersion cluster. **(A)** Division of word epochs as a function of surprisal and dispersion. (B) Power event-related potentials (ERPs), baseline corrected, of the different quadrants in the Dispersion cluster. (C-D) Strength cluster. (C) Division of word epochs as a function of surprisal and strength. (D) Power ERPs, baseline corrected, of the different quadrants in the Strength cluster. Colored lines correspond to statistically relevant activity compared to a *−*1000 ms to *−*500 ms baseline. Transparent blue/red vertical areas correspond to the time window of maximal uncertainty-surprisal synergy, as defined by the PID analysis (Fig.3J). Blue/red arrows represent significant increases in power within the synergy window between two adjacent quad-rants. (E,F) Effect of the combination of uncertainty and surprisal, compared to an additive model of these features (green), for the dispersion ((E)) and strength ((F)) clusters.

In the dispersion cluster, all categories elicited significant responses (*p <* 0.01, FDR corrected), with activity from ∼ 50 ms and a sharp peak at 300 ms (Fig.4B). This population thus responds broadly to words, consistent with a sensory processing role. Dispersion influenced responses only when words were not surprising, whereas surprisal contributed regardless of dispersion (*p <* 0.05, FDR-corrected). The response in the high-dispersion + high-surprisal quadrant was weaker than predicted by the sum of single-dimension effects (two-tailed t-test, *t* = −2.76*, df* = 247*, p* = 0.005; Fig.4E), a sub-additive interaction which indicates that uncertainty dampens surprising inputs, in line with a Bayesian predictive framework (Friston, 2010).

In sharp contrast, the strength cluster responded exclusively to highly surprising words (*p <* 0.05, FDR-corrected), regardless of prediction strength, with late sustained activity from 250–500 ms, akin to an N400 (Sanders and Neville, 2003) (Fig.4D). Additivity analysis showed a supra-additive interaction (*t* = 2.08*, dof* = 163*, p* = 0.03; Fig.4F), i.e. uncertainty amplifies the response to surprising inputs, consistent with an information-seeking framework (Roesch et al., 2012).

Together, these results highlight distinct functional roles for dispersion and strength. Dispersion dampens surprising inputs in lower-level sensory populations, optimizing sensory coding, whereas strength amplifies responses to surprising words in higher-order lexical populations, reflecting statistical rather than sensory processing and promoting exploratory processing.

### A partially hierarchical organization of dispersion and strength clusters

Predictive coding theory posits that higher-level errors arise from lower-level errors propagation, implying that responses in the strength cluster should follow those in the dispersion cluster. Instead, we observed substantial temporal overlap between the two (Fig.4B,D), which deviates from this account.

To assess information flow directly, we applied feature-specific information transfer (FIT) (Celotto et al., 2023), an information-theoretic measure of directed commu-nication between neural sites (Fig.5A). FIT computed for dispersion, strength, and surprisal showed similar patterns due to their shared information, with the clear-est effects for dispersion (Fig.11). This analysis revealed a significant bidirectional exchange between the two clusters (permutation cluster, *p <* 0.001; Fig.5B). Before word onset, information flowed top-down from the higher-level strength cluster to the sensory dispersion cluster, while around 300 ms it flowed bottom-up from the disper-sion to the strength cluster. The average delay of information transfer between the two clusters (∼ 260 ms) suggests indirect rather than direct connections.

**Fig. 5.**
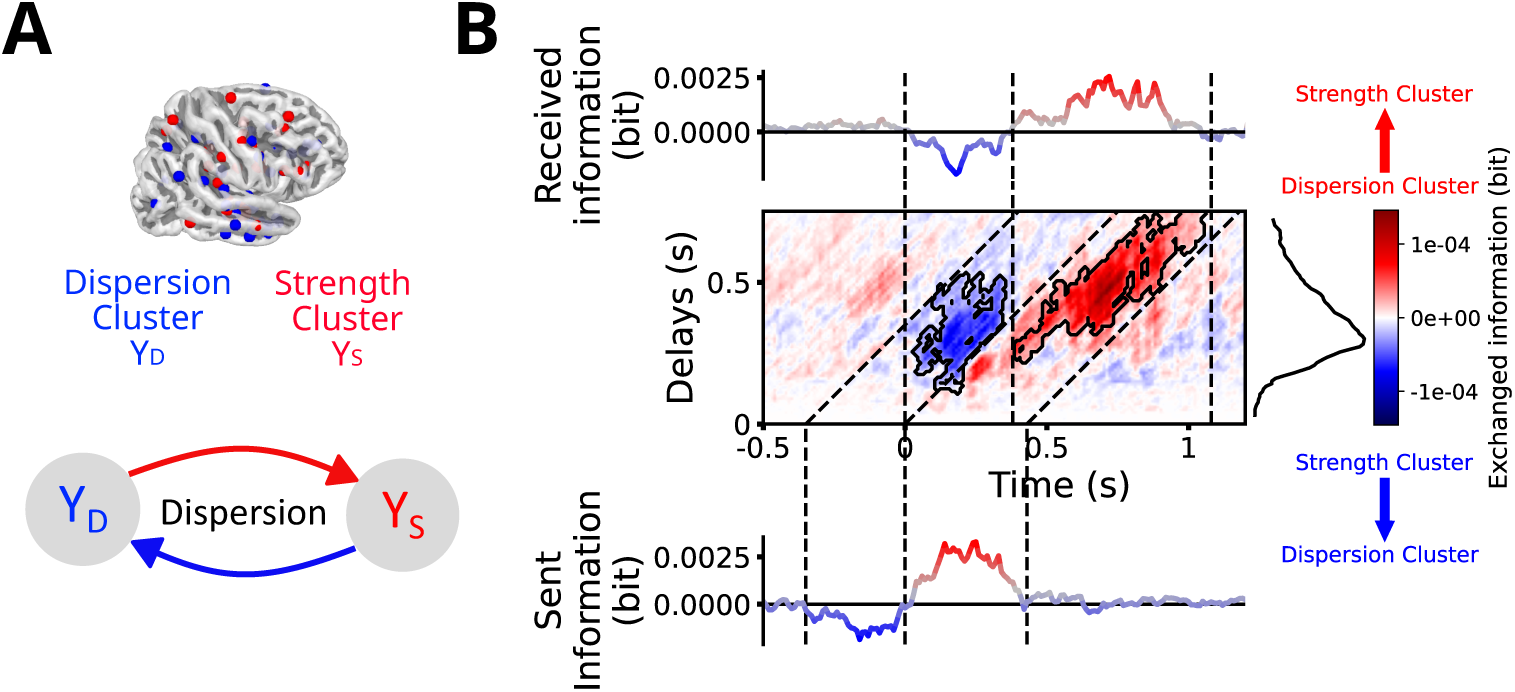
Feature-specific information transfer (FIT) relative to dispersion between the dispersion and strength clusters. **(A)** Dispersion FIT framework: for each pair of channels across clusters within the same subject, the net information transfer of dispersion is computed. **(B)** Time-delay FIT map. The transferred information was summed across delays to compute the received information (top), and summed across diagonals to compute the sent information (bottom). The absolute value of information was averaged across times to compute the average delay between sender and receiver (right). Statistical significance in the FIT map is displayed using contours. Negative values (blue) correspond to a transfer from the strength cluster to the dispersion one, while positive values (red) correspond to a transfer from the dispersion cluster to the strength one. The information was first sent from the strength cluster around *−*150 ms pre-word to the dispersion cluster around 100 ms post-onset. It was then sent back from the dispersion cluster between 200 ms and 350 ms post-word onset to the strength cluster from 600 ms to 800 ms post-word onset.

These results demonstrate a bidirectional exchange of information between the two clusters (Heilbron et al., 2022). Yet the latency pattern (Fig.5B) departs from classical predictive coding: the strength cluster generates prediction errors independently, before bottom-up signals from the dispersion cluster arrive. Predictive processing therefore arises not from a single hierarchical chain but from the interplay of partially independent local and global circuits (Brodbeck et al., 2022).

## Discussion

While Shannon entropy is the standard metric for quantifying uncertainty in both computational and cognitive neurosciences, our findings show that the brain does not represent lexical uncertainty in this form. Rather, it relies on two complementary measures: the probability of the most likely event (strength) and the number of plausible events (dispersion). Hence, when faced with a large lexical space, the brain does not encode the full discrete probability distribution over lexical candidates but approximates it with two continuous summary variables. These uncertainty measures operate at different stages of the speech processing hierarchy, with dispersion associated to sensory prediction and Bayesian optimization, and strength associated to lexical processing and information gain. Each stage thus relies on summary statistics aligned with its computational goal. Finally, our findings challenge a strictly hierarchical view of predictive coding, pointing instead to a distributed architecture in which local computations drive predictive processing while being modulated by error propagation. When multiple alternatives are possible, the brain relies on summary statistics rather than on the whole probability distribution, which can lead to suboptimal behavior (Churchland et al., 2008; Yeon and Rahnev, 2020; Ma and Jazayeri, 2014). The choice of the summary statistics is non-trivial for complex distributions and tasks (Daikoku, 2018), such as language processing. Our results demonstrate that during speech comprehension, lexical uncertainty is represented as the two extremes of this family: the uncertainty relative to the number of plausible alternatives (*α* → 0; dispersion) and the uncertainty of the most probable word (*α* → ∞; strength). Dispersion converges to the Hartley entropy *H*_0_(*X*) (Wehrl, 1990) and therefore captures neighborhood size or lexical cohort density (Kowialiewski and Majerus, 2019). (In practice, we computed it as *H*_0.01_(*X*) to account for the non-zero probabilities of language models.) In contrast, strength corresponds to the Bayes error (also coined error entropy or min-entropy (Fano and Hawkins, 1961)), relies fully on the notion of probability and focuses on a single event, making it particularly relevant for prediction tasks (Yeon and Rahnev, 2020). Critically, the neural representation of strength is limited to the most likely upcoming word, with no evidence of significant encoding of the probability of the second most likely word (Fig.10A,B,E-L). Thus, we propose that in a large lexical space, the brain does not encode the full discrete distribution of word-level probabilities but instead approximates it with two continuous summary statistics: dispersion and strength. These are analogous to the variance and the mode probability of a continuous distribution, consistent with theories that neural codes represent uncertainty via low-dimensional parameters rather than full distributions Knill and Pouget (2004); Pouget et al. (2013); Ma and Jazayeri (2014). In this view, predictive processing reduces the combinatorial complexity of lexical probability space by mapping it onto continuous approximations. What is considered high-dimensional for human brains is likely task-dependent, and remains for future work to be addressed. Dispersion and strength are processed by distinct neural populations at different stages of the speech processing hierarchy: dispersion in early sensory regions and strength in higher-level lexical areas. Dispersion shapes sensory predictions, notably pre-word activity, likely reflecting anticipatory representations of plausible auditory inputs (Kok et al., 2017). This anticipatory activity continues and peaks around 100 ms, a signature of word-onset responses (Sanders et al., 2002; Brodbeck et al., 2018) and early sensory processing (Näätänen and Picton, 1987). We thus posit that this peak reflects the first comparison between the expectations set by anticipatory representations and the actual input. Furthermore the later response around 300 ms can be interpreted as reflecting early, pre-lexical phonetic mismatch (Connolly et al., 2001; Newman and Connolly, 2009). The latter is supra-additively modulated by joint dispersion and surprisal, consistent with Bayesian accounts of sensory processing, in which uncertainty dampens surprising inputs (Friston, 2010; Meyniel, 2020). In sharp contrast, the strength population displays no sensory activity, only a late, sustained response to surprising words, characteristic of higher-order lexical processing (Tyler et al., 2005; Frank et al., 2015; Huth et al., 2016) and akin to a N400 (Sanders and Neville, 2003). Contrary to dispersion, strength interacts supra-additively with surprisal: the lack of a strong candidate for the next word amplifies the neural response to unexpected words, thus promoting information-seeking computations (Roesch et al., 2012; Richter et al., 2018; Song et al., 2024). This suggests that the lexical stage commits to a single best prediction, directly compared to incoming input. Whether surprisal or another dissimilarity measure (semantic distance for example) is the most fitted metric to characterize the input at this processing stage still needs to be answered. Regardless, dispersion and strength support opposing neural functions, across different timescales and neural populations. These findings are consistent with the opposing process theory (Press et al., 2020), which posits that both Bayesian (sensory optimization) and cancellation (information-seeking) mechanisms contribute to perceptual inference, but operate at different timescales. The speech processing hierarchy thus relies on complementary summary statistics specific to the computational goal of each stage. Yet, both populations show temporally overlapping responses to surprising words, indicating that higher-level surprisal is not solely inherited from sensory errors. Instead, sensory and lexical circuits operate largely in parallel, while also exchanging top-down predictions and bottom-up errors (Brodbeck et al., 2022). Thus, predictive coding in language relies on local predictive circuits supported by inter-areal communication, rather than from a sequential error-propagation chain.

Although our study focused on lexical predictions, language processing operates across multiple representational levels. Different linguistic units (phrases, syllables, phonemes) likely present different probability distributions, neural functions and populations, and therefore distinct entropy profiles (Donhauser and Baillet, 2020; Gibson et al., 2019). Dissociating their respective contributions is a complex task and will be the subject of future inquiries. While our findings are specific to speech, the parameterization of uncertainty into complementary measures can generalize to other domains of prediction. We thus advocate for a broader use of generalized entropy families (Crupi et al., 2018; Sharma and Mittal, 1975), as they offer a simple and powerful way to capture distinct forms of uncertainty and to disambiguate the neural computations that underlie complex cognition.

## Materials and methods

### Data and code availability

The raw data investigated in the current manuscript is privileged patient data. Therefore the conditions of our ethics approval do not permit public archiving of anonymised data. Readers seeking access to the data should contact Dr. Daniele Schön. Access will be granted to named individuals in accordance with ethical procedures governing the reuse of clinical data, including completion of a formal data sharing agreement. Data analyses were performed using custom scripts in Python, which are available on GitHub : https://github.com/phg17/Renyi-entropy.

### Participants

Stereotactic electroencephalography (sEEG) recordings were obtained from 33 patients (18 women; mean age, 30 years; range, 8 to 54) with refractory epilepsy using intracerebral electrodes implanted as part of the standard presurgical evaluation process. Electrode placements were based solely on clinical requirements. All patients were French native speakers. Neuropsychological assessments carried out before sEEG recordings indicated that they all had intact language functions and met the criteria for normal hearing. None of them had their epileptogenic zone including the auditory areas as identified by experienced epileptologists. Recordings took place at the Hôpital de La Timone (Marseille, France). All patients gave informed consent, and the experiment reported here was approved by the Assistance Publique – Hôpitaux de Marseille (health data access portal registration number PADS E2YSEB).

### sEEG recordings

The sEEG signal was recorded using depth electrodes shafts of 0.8-mm diameter containing 10 to 15 electrode contacts (Dixi Medical or Alcis, Besançon, France). The contacts were 2 mm long and were spaced from each other by 1.5 mm. Data were recorded using a 256 -channels amplifier (Brain Products), sampled at 1000 Hz and high-pass filtered at 0.016 Hz inside a soundproof insulated Faraday cage. The recording reference and ground were chosen by the clinical staff as two consecutive sEEG contacts on the same shaft both located in the white matter and/or at distance from any epileptic activity. Neural recordings were performed between 3 and 10 days after the implantation procedure. No sedation or analgesics drugs were used, and antiepileptic drugs were partially or completely withdrawn. Recordings were always acquired more than 4 hours after the last seizure. Patients were included in the study if their implantation map covered at least partially the Heschl’s gyrus (left or right). 31 of 33 patients were implanted bilaterally, yielding a total of 544 electrodes and 5194 contacts (Fig.11A). Patients were implanted with an average of 16 (range, 11 to 19) depth electrodes.

### Preprocessing

To increase spatial sensitivity and reduce passive volume conduction from neighboring regions (Mercier et al., 2017, 2022), the signal was offline re-referenced using bipolar montage. That is, for a pair of adjacent electrode contacts, the referencing led to a virtual channel located at the midpoint locations of the original contacts. To precisely localize the channels, a procedure similar to the one used in the iELVis toolbox and in the FieldTrip toolbox was applied (Groppe et al., 2017; Stolk et al., 2018). First, we manually identified the location of each channel centroid on the post-implant CT scan using the Gardel software (Villalon et al., 2018). Second, we performed volumetric segmentation and cortical reconstruction on the pre-implant MRI with the Freesurfer image analysis suite (documented and freely available for download online at http://surfer.nmr.mgh.harvard.edu/). This segmentation of the pre-implant MRI with SPM12 provides us with both the tissue probability maps (i.e. gray, white, and cerebrospinal fluid [CSF] probabilities) and the indexed-binary representations (i.e. either gray, white, CSF, bone, or soft tissues). This information allowed us to reject electrodes not located in the brain. Third, the post-implant CT scan was coregistered to the pre-implant MRI via a rigid affine transformation and the pre-implant MRI was registered to MNI152 space, via a linear and a non-linear transformation from SPM12 methods (Penny et al., 2011), through the FieldTrip toolbox (Oostenveld et al., 2011). Fourth, applying the corresponding transformations, we mapped channel locations to the pre-implant MRI brain that was labeled using the volume-based Human Brainnetome Atlas (Fan et al., 2016).

Based on the brain segmentation performed using SPM12 methods through the FieldTrip toolbox, bipolar channels located outside of the brain were removed from the data (3%). The remaining data was then bandpass filtered between 0.1 and 250 Hz, and, following a visual inspection of the power spectral density profile of the data, when necessary, we additionally applied a notch filter at 50 Hz and harmonics up to 200 Hz to remove power line artifacts. Finally, the data were downsampled to 100 Hz.

### Experimental Design

The experiment consisted in a single session in which patients passively listened to ∼ 10 min of storytelling (577 s), La sorcìere de la rue Mouffetard (Gripari and Dahl, 1994). The recording session was conducted in the Faraday cage (N=24) or in the bedroom (N=10).

In the Faraday cage, a sound Blaster X-Fi Xtreme Audio, an amplifier Yamaha P2040 and Yamaha loudspeakers (NS 10M) were used for sound presentation. In the bedroom, stimuli were presented using a Sennheiser HD 25 headphone set. Sound stimuli were presented at 44.1 kHz sample rate and 16 bits resolution. Speech and music excerpts were presented at 75 dBA.

### Computations of predictions via a Large Language Model

The neural recordings were compared with the predictions of a pretrained large language model (LLM) (Radford et al., 2019) in response to the same story as presented to subjects. Because the stories were in French, we used the Camembert model (Martin et al., 2019), a performant language model based on RoBERTa (Liu et al., 2019) and fined-tuned for French. We used the pretrained models from Huggingface. For each word of the story, the model used all previous words as context to make predictions about the next one. These predictions take the form of a probability distribution over the 32k subword tokens.

### Computation of surprisal and uncertainty

Given the distribution of possible tokens, we computed the surprisal of the first token of each observed word as:

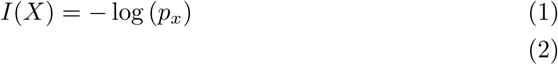

with *p_x_* the probability of the observed token.

Traditionally, uncertainty is computed as the Shannon entropy over the distribution of possible events (tokens) i.e. the expected value of surprisal:

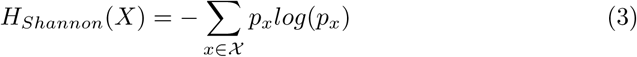

with *χ* the distribution of tokens. While this formulation captures the variety of distribution shapes for two events (Fig.6B), it becomes ambiguous for more events (Fig.6E). Here, we sought to refine the formulation of uncertainty, using a generalization of the Shannon entropy: the Rényi entropy ((Rényi, 1961)). The latter relies on a weaker postulates about the additive property of entropy. This allows to extend the Shannon entropy to a family of quantities, characterized by their Rényi order *α*, as:

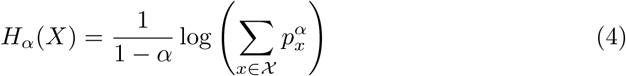

This newly defined quantity allows to give different weights to unlikely events (Fig.6I), as they lose importance with higher *α*. As such, when *α* → 0, all plausible events (events with non-null probability) are considered equally, this case is referred to as the Hartley entropy or dispersion. On the contrary, when *α* → +∞, only the probability of the most likely event is considered. Hence, every intermediary value of *α* represents different trade-offs between dispersion (the number of plausible events) and strength (the probability of the most likely event) (Fig.6K). This quantity further covers the classical Shannon entropy as *H_α_*_→1_ = *H_Shannon_*. Uncertainty is thus not defined by a single quantity, but rather by an entropy profile, which allows to distinguish between different distribution shapes (Fig.6C,F,J). Hence, given the probability distribution of all possible tokens at every word onset, the uncertainty was computed as the Rényi entropy for 20 values of *α*: 0.01, 0.05, 0.1, 0.2, 0.3, 0.4, 0.5, 0.6, 0.7, 0.8, 0.9, 1, 1.1, 1.2, 1.3, 1.4, 1.5, 2, 5 and 10. We have approximated Dispersion by *α* = 0.01 since the output of the language never outputs exactly null probabilities and the strictly defined Hartley entropy is thus be constant across the dataset (Klir, 2006). Critically, *H_α_*(*X*) monotonously decreases with *α*, meaning that entropies associated to different *α* cannot be compared directly. For the same *α* however, we may compare the entropies associated to different words, specifically their ranks within the corpus (Fig.14A-C) or z-score (Fig.8C, Fig.7E-H), consistent with our rank-based approach. Normalization allows us to verify that, during a sentence, all forms of uncertainty decrease (Fig.8A,B).

### Gaussian Copula Mutual Information

To quantify the statistical dependency between neural data and each of our 20 measures of uncertainty, we used an information-based measure: the mutual information (MI), defined as:

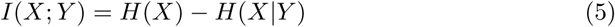

where *X* and *Y* represent the broadband neural data epoched over words and the uncertainties at each of these, with *H*(*X*) and *H*(*Y*) their respective entropy. *H*(*X*|*Y*) is the conditional entropy of X given Y. More specifically, we adopted the Gaussian Copula Mutual Information (GCMI) (Ince et al., 2017), which does not require binning to compute MI. It is a robust rank-based approach that allows to detect non-linear monotonic relation between variables and is a lower-bound approximate estimator of MI. This method is well-adapted to our informational features as the neural coding of probability is often non-linear (Foucault et al., 2024). Before using this framework, we first transformed all variables (features and neural data) into Gaussian variables using the inverse normal transformation. The MI was then computed as:

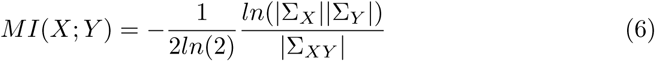

with Σ*_X_*, Σ*_Y_* and Σ*_XY_* respectively being the covariance matrices of X, Y and of the joint variable (*X, Y*). The GCMI was computed at each time-point of the epochs, for every channel. We thus obtained a time-series for each channel and for each uncertainty. All computations were performed using the frites toolbox (Combrisson et al., 2022)

We thus obtained timecourses of mutual information for each channel and each value of *α*. From these, we extracted the value at the peak of the averaged timecourse for each channel and value of alpha, resulting in a single value per channel and per Rényi order. This allowed us to plot the entropy profile (Fig1C, Fig2B,E).

### Non-negative Matrix Factorization

To characterize the heterogeneity in preferred entropy measures across channels, we applied non-negative matrix factorization (NMF) as a soft clustering method, appro-priate given the strictly positive nature of mutual information (MI) (Yu et al., 2024; Lee and Seung, 1999). The NMF iteratively approximates a nonnegative data matrix **X** ∈ *R^N,p^*by:

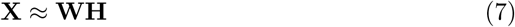

where **W** ∈ *R^N,r^*and **H** ∈ *R^r,p^*are nonnegative matrices respectively referred to as the basis matrix and the coefficient matrix, with *r* ≤ *max*(*N, p*) the number of compo-nents. The coefficient matrix **H** represents how strongly a data point belongs to each of the component and can therefore be interpreted as a membership matrix (Li and Ding (2018); Fig.10C). Using a threshold on the latter, we can obtain a soft clustering of the data. For each time series previously obtained, we computed the average MI timecourse (from word onset to the 95% word offset), thus obtaining 20 strictly positive data points per channel, on which the NMF was applied. Hyperparameters (number of clus-ters, regularization parameters) were optimized using the silhouette score(Shahapure and Nicholas, 2020). Channels with a soft membership below 0.65 were excluded to retain only those with strong cluster membership. We further tested different cluster threshold (0.5 and 0.8) to ensure the consistency of results (Fig.13A-D).

### Partial Information Decomposition (PID)

The goal of studying different definition of entropy is to be able to identify the one that best explain the neural data. However, this task is non-trivial as different forms of entropy are highly correlated (Fig.7A-C). Therefore, it is necessary to be able to dissociate the unique and redundant information they contain about the neural data. Moreover, one essential aspect of predictive coding is the interaction between prediction(uncertainty) and the observed signal (the surprisal), which also display shared information (Fig.7D). To address these two questions, we employed the Par-tial Information Decomposition (PID) (Williams and Beer, 2010; Wibral et al., 2015) framework. The latter decompose the joint mutual information that two variables *X*_1_ and *X*_2_ hold about a third variable *Y*, into 4 information atoms: the atoms uniquely attributed to respectively *X*_1_ and *X*_2_, the atom redundantly attributed to both and the atom synergistically created when observed together:

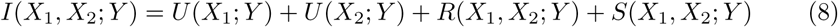

To calculate this decomposition, we first computed the redundancy measure as the minimum of the information provided by each individual feature about the target ((Barrett, 2015)):

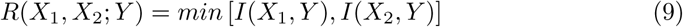

The unique information terms can then be computed algebraically as:

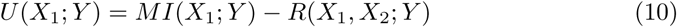

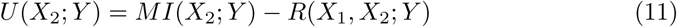

And finally synergy is given by:

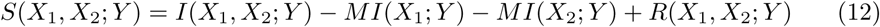

This approach is most commonly used to study how the combination of different neural signal informs about a single feature(Combrisson et al., 2024; Greco et al., 2024). In our case, it is more fit to quantify the contribution of different features on a single channel (Daube et al., 2019). Hence, the unique information allows to determine the specificity to which a channel respond to one feature over another, while redundancy and synergy informs us about the interactions between them (Ince et al., 2017; Kay and Ince, 2018). All computations were performed using the frites toolbox (Combrisson et al., 2022).

For each component of the PID and each cluster, we used the information from −500 ms to −250 ms pre-word as a baseline to establish statistical relevancy of the information between −250 ms and 1250 ms relative to word onset.

### Feature-specific Information Transfer (FIT)

To characterize the transfer of information between two channels, *Y*_1_ and *Y*_2_, about a feature, *X*, it is necessary to combine an information theory approach with Granger causality principle. To achieve that, we used the feature-specific information trans-fer (FIT). This method identifies the information about a feature, *X* a channel, *Y*_2*,pres*_ has in common with the past of another channel, *Y*_1*,past*_, while controlling for its own past, *Y*_2*,past*_, (Celotto et al., 2023; Lemke et al., 2024). It relies on the PID introduced earlier and can be formulated as follow:

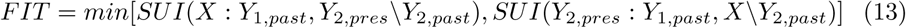

where *SUI*(*X* : *Y*_1*,past*_*, Y*_2*,pres*_\*Y*_2*,past*_) is the information shared by *Y*_1*,past*_ and *Y*_2*,present*_ about *X* that is not contained in *Y*_2*,past*_. *SUI*(*Y*_2*,pres*_ : *Y*_1*,past*_*, X*\*Y*_2*,past*_) is the information shared by *Y*_2*,pres*_ and *X* about *Y*_1*,past*_ that is not contained in *Y*_1*,past*_. As such, *FIT* (*Y*_1_ → *Y*_2_; *X*) quantifies the amount of information about the feature, *X*, transferred from *Y*_1_ to *Y*_2_.

Here the FIT was used to assess the directionality of the information transfer between our two clusters, relative to our different information features. To do so, we computed both *FIT* (*Y*_1_ → *Y*_2_; *X*) and *FIT* (*Y*_2_ → *Y*_1_; *X*) and subtracted one from the other for each lag and channel. This resulted in a time-delay matrix, containing the amount of information being asymmetrically transmitted between the two clusters at a certain time and from a certain delay in the past. Negative and positive values thus signifies opposite direction for the information transfer i.e. the preferred direction of information transfer (Bastos et al., 2015). This metric was computed over all possible pairs of channels from both clusters within each subject separately, resulting in 956 pairs. The FIT was computed for delays going up to 1 second. Statistical significance of time-delay regions was assessed using permutation clusters with 1000 permutations across pairs of channels. By averaging the matrix across delays, we computed the timecourse of average received information. Then, by averaging data along time-delay diagonal, we obtained the timecourse of average sent information. Finally, by averaging along timepoints, we could obtain the average delay of communication between clusters. Contrary to the PID, the FIT does not allow to dissociate the contribution of different features, the patterns of information transfer between redundant features are therefore similar (Fig.11B), limiting the possible interpretations.

### Orthogonalisation of Surprisal and Uncertainty

While information-based metrics allow to detect and quantify non-linear statistical dependencies between features and neural data, it does not allow to characterize the measured effect. Therefore, in order to understand the interaction between uncertainty and surprisal within each cluster, we adopted a data-partition approach. We separated the words in our corpus according to their surprisal and the uncertainty of their associated context. This procedure was done separately for each form of uncertainty revealed by previous analysis: the Rényi entropies for *α* = 0.01 and *α* = 10. First we took out outliers in terms of word duration, cutting words shorter than 100 ms and longer than 700 ms. We then represented each word within the corpus as a point in a 2D space, with the x-axis representing uncertainty and the y-axis surprisal. Then, within that space, we selected the points that fell within the first and last 30% for the surprisal and within the within the first and last 45% for the uncertainty. These values were chosen as to keep at least 10% of the total number of words within each of the quarters we thus defined (Fig.4A,C). Then using a procedural methods, we extracted from each square, 5% of the total number of words as to minimize the Bayes factor of the distribution of the data projected on each axis, for their respective neighboring quarter (Kass and Raftery, 1995). This resulted in 4 orthogonal quarters of words: low-uncertainty + low-surprisal (bottom left), low-uncertain + high surprisal (top left), high-uncertainty + low-surprisal (bottom right), high-uncertain + high surprisal, for both dispersion and strength (all *bf <* 0.18, Fig.12A-D).

### ERP power analysis

We computed the event-related potential (ERP) power for each of the 4 categories, as well for all words (Fig.10D), relative to the cluster and uncertainty of interest, using the −1000 ms to −500 ms activity pre-word onset as a baseline to assess statistical relevance. The ERP power was computed since the use of a bipolar montage prevents polarity from being interpretable. Comparison between quadrants was done based on a 100 ms window around the peaks of synergy between uncertainty and surprisal highlighted by the PID: 300 ms post-onset for both the dispersion and strength cluster. Effects of the different features were computed as the difference between the synergy region in a quadrant compared to an orthogonal quadrant (Fig.4B,D, colored arrows).

The joint effect of dispersion and surprisal was characterized by first computing for each channel an additive model as the sum of the effect of surprisal in the absence of dispersion (from lower-left to upper-left), and of dispersion in the absence of surprisal (from lower-left to lower-right). This additive model was then compared to the combined effect of surprisal and dispersion (from lower-left to upper-right) to establish additivity, sub-additivity or supra-additivity using a two-tailed t-test.

### Spatial representation

Because of the sparsity of intracranial implantation, spatial representations do not generalize well. Here we ran a procedure, akin to a searchlight procedure, in order to delete spatially isolated channels, or groups of channels driven by a single subject. When spatially representing our data(Fig1E, Fig2G), we first considered the neighborhood of each point as a 10 mm sphere around them, and only plotted channels that contained a channel from a different subject within their neighborhood.

Moreover, in order to compare the spatial representation of each cluster of activity, we reduced their position within the 3D space as their distance to the primary auditory cortex, as te1.1 (Kiviniemi et al., 2009; Anderson et al., 2011; Norman-Haignere et al., 2024). Their distances could then be compared using a t-test.

## Acknowledgments

This work was funded by the European Union (ERC, SPEEDY, ERC-CoG-101043344), Fondation Pour l’Audition (FPA RD-2022-09), France 2030 (ANR-16-CONV-0002) and the Excellence Initiative of Aix-Marseille University (A*MIDEX AMX-19-IET-004). We would like to thank A. Brovelli, JR. King, V. Wyart, D. Schön and the members of the D-cap team at INS for thoughtful comments on the study. We would like to express our gratitude to the patients and their families for their time and commitment. We also acknowledge the wonderful work done by the nurses and doctors at the epilepsy unit. Last, we wish to thank S. Guilleminot.

## Author information

These authors contributed equally: Pierre Guilleminot, Benjamin Morillon.

## Contributions

BM acquired the data, PHG processed the data, PHG and BM interpreted the data, wrote and revised the manuscript. Both authors approved the final manuscript.

## Corresponding author

Correspondence to Pierre Guilleminot.

## Ethics declarations

The authors declare no competing interests. The study was approved by the Assistance Publique – Hôpitaux de Marseille (health data access portal registration number PADS E2YSEB).

## Supplementary information

**Fig. 6.**
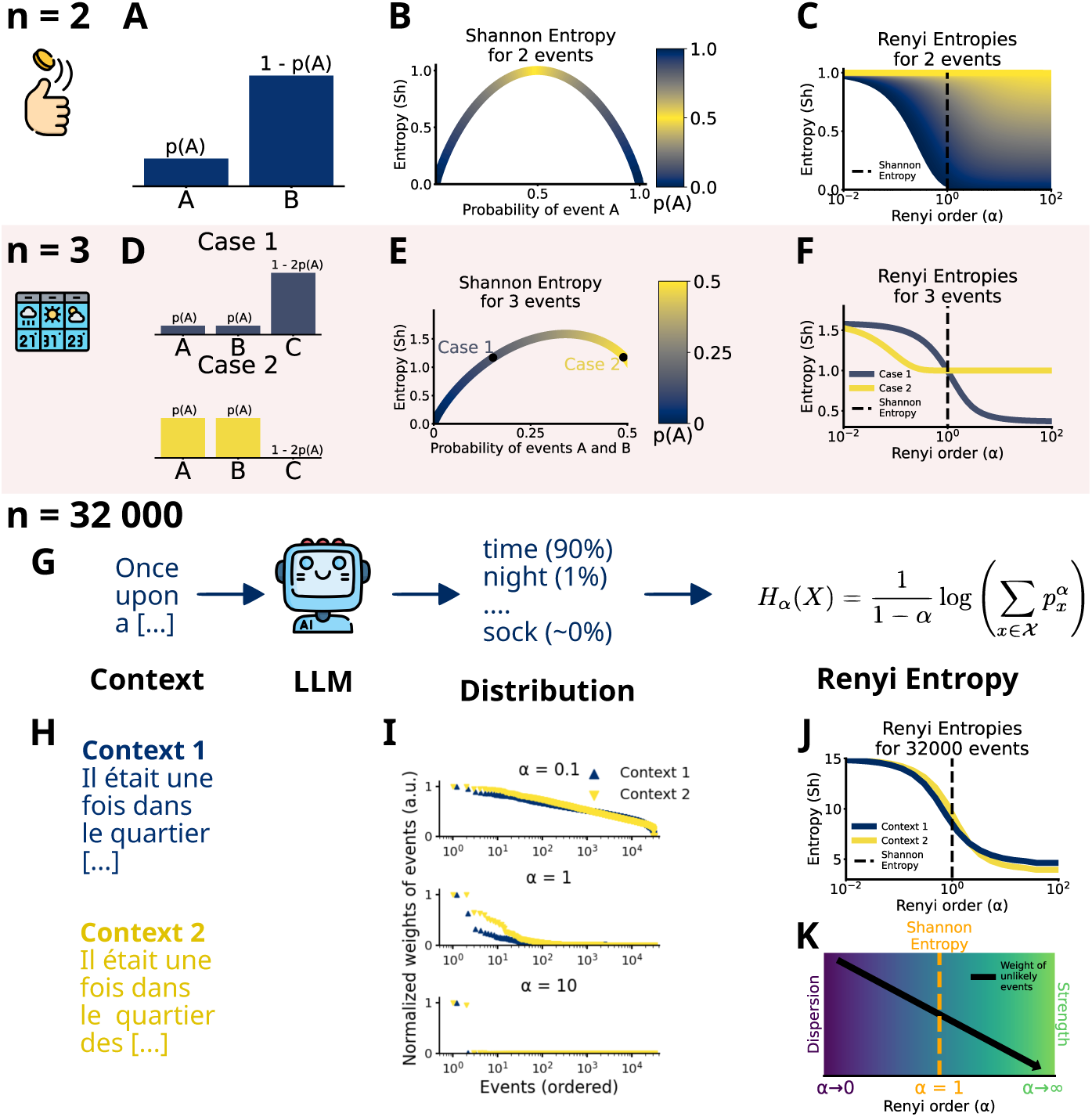
Extended Data 1. **(A)** A 2-event distribution, fully described by p(A). **(B)** Shannon entropy for *n* = 2 as a function of p(A). **(C)** Rényi entropy as a function of the Rényi order *α*, for different p(A) (color gradient). *α* = 1 corresponds to Shannon entropy (dotted line). **(D)** Two cases of 3-event distributions with two equiprobable events, fully described by p(A). **(E)** Shannon entropy for *n* = 3 as a function of p(A) (= p(B), color gradient). **(F)** Rényi entropy as a function of *α*, for cases 1 and 2, which have the same Shannon entropy (dotted line). **(G)** In Large Language Models (LLMs), contextual probabilities are assigned to all possible words (*n* = 32000 events). **(H)** Two contexts example. **(I)** Normalized weights of events for different *α*. **(J)** Rényi entropy as a function of *α*, for contexts 1 and 2. **(K)** The entropies describing the most likely event (*α → ∞*, green) and all non-null events (*α →* 0, night blue) are respectively referred to as the Prediction Strength (Str) and the Distribution Dispersion (Dis).

**Fig. 7.**
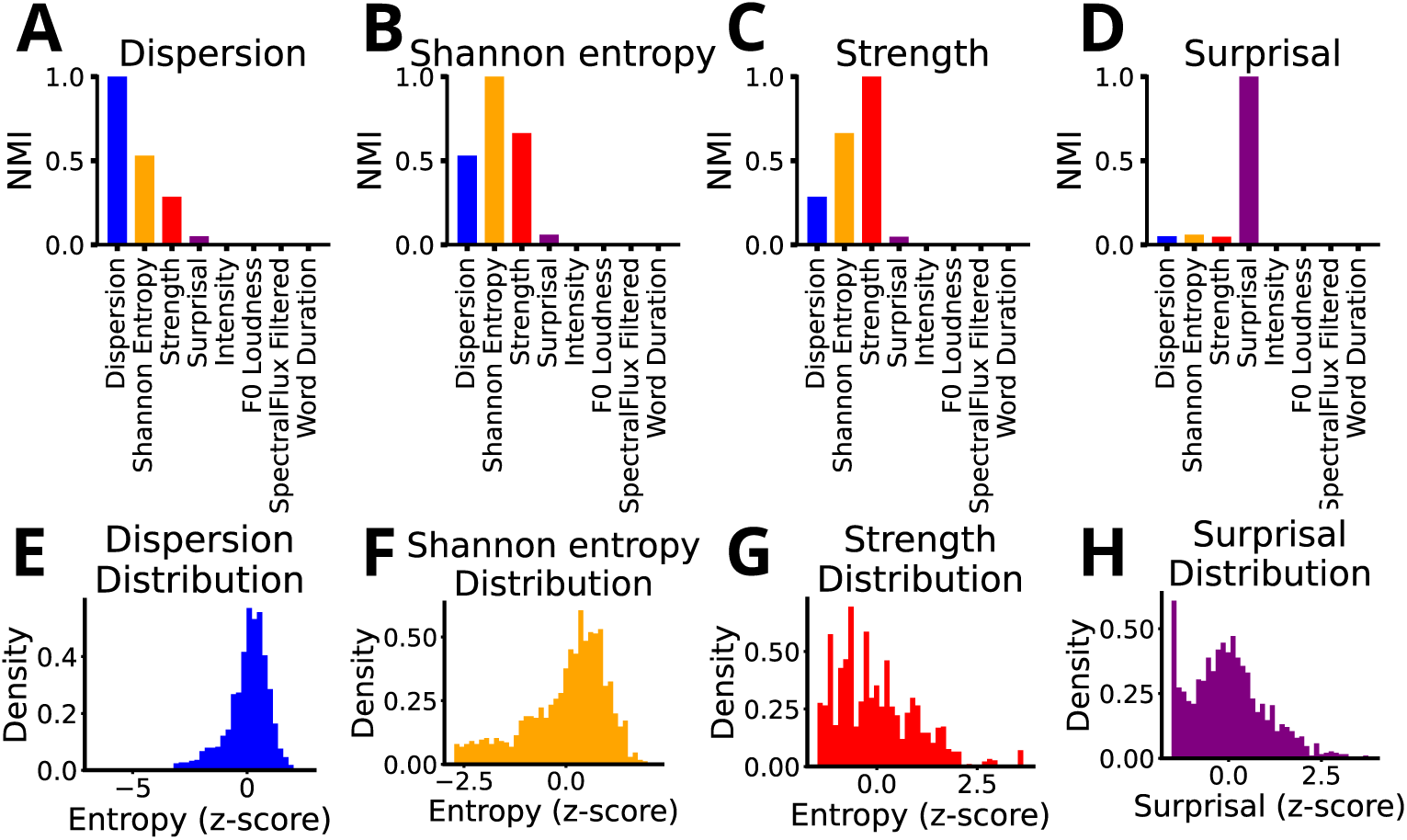
Extended Data 2. **(A-D)** Normalized mutual information (NMI) between speech features and dispersion **(A)**, Shannon entropy **(B)**, strength **(C)** and surprisal **(D)**. **(E-H)** Distribution of the values of dispersion **(E)**, Shannon entropy **(F)**, strength **(G)** and surprisal **(H)**

**Fig. 8.**
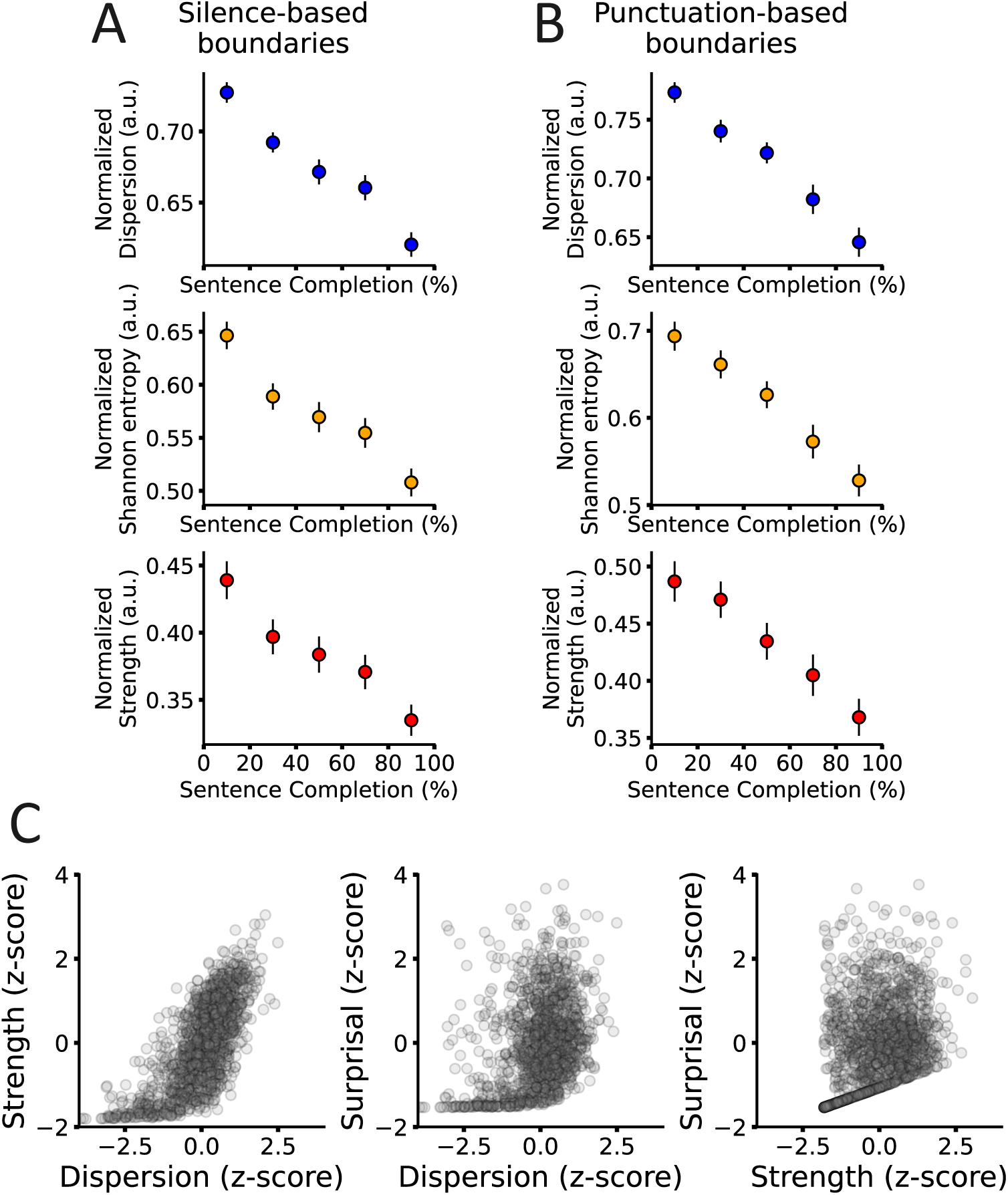
Extended Data 3. **(A,B)** Values of dispersion (blue), Shannon entropy (orange) and strength(red) at a function of the position within a sentence. Positions were normalized according to sentence length and grouped by bins of 20%. All measures of uncertainty decrease throughout sen-tences. Sentences were defined according to either punctuation within the text **(A)** or silences in the audio **(B)**. **(C)** Scatterplot of strength vs dispersion, surprisal vs dispersion and surprisal vs strength.

**Fig. 9.**
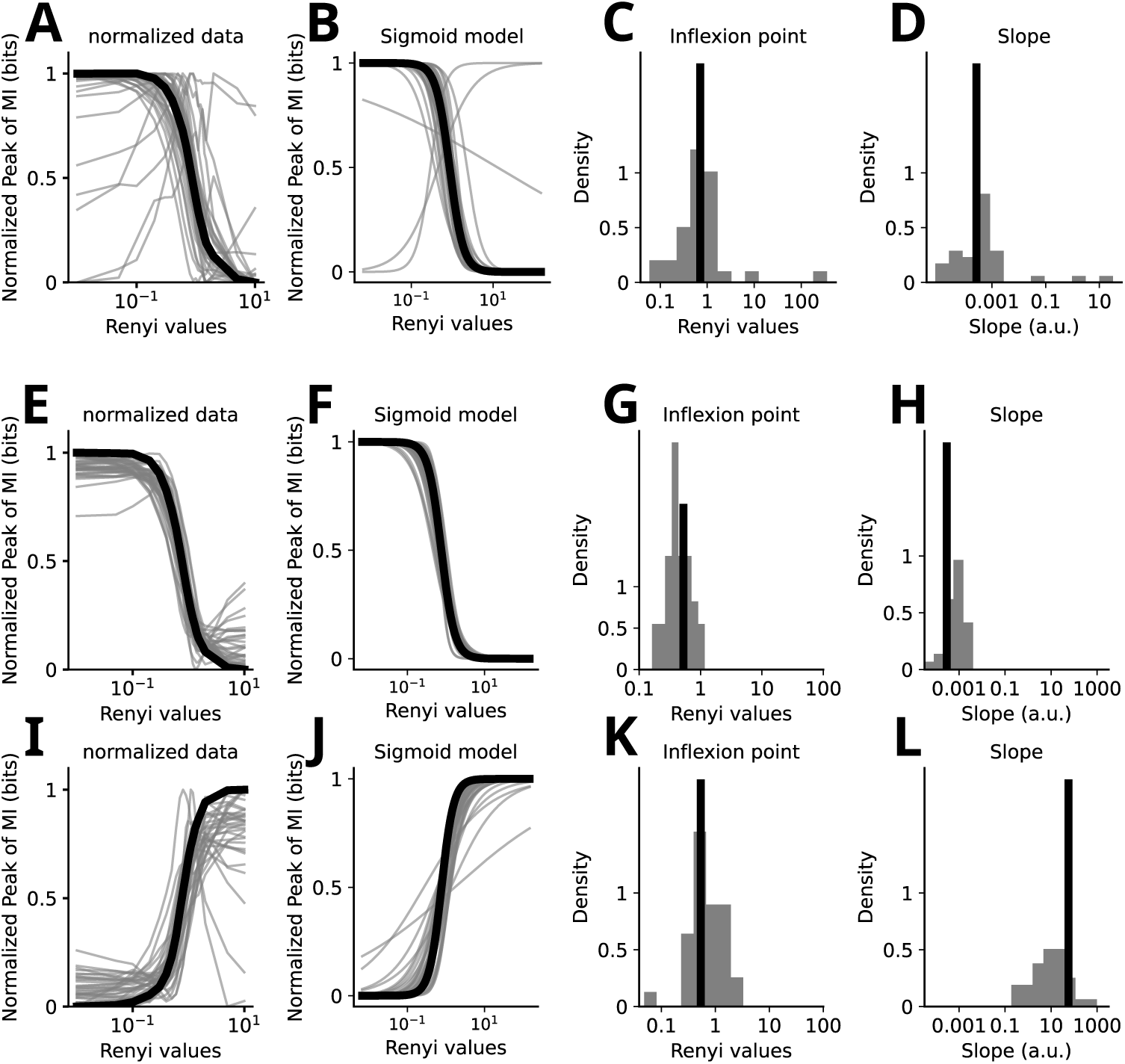
Extended Data 4 (A-D) Individual entropy profiles for the average mutual information. Entropy profiles were averaged by subject and normalized between 0 and 1 **(A)**, which resulted in one profile per subject (grey lines) and an average profile (black line). **(B)** Using least-square optimization via a trust region reflective algorithm, we fitted a sigmoid to both average and individual profiles, resulting in a sigmoidal model per subject (grey lines) and a average one (black line). **(C,D)** Distribu-tion of inflection points **(C)** and slope **(D)** for each individual models. This procedure was repeated identically using only channels in the dispersion cluster **(E-H)** and in the strength cluster **(I-L)**. This shows that the cluster separation is consistent across subjects.

**Fig. 10.**
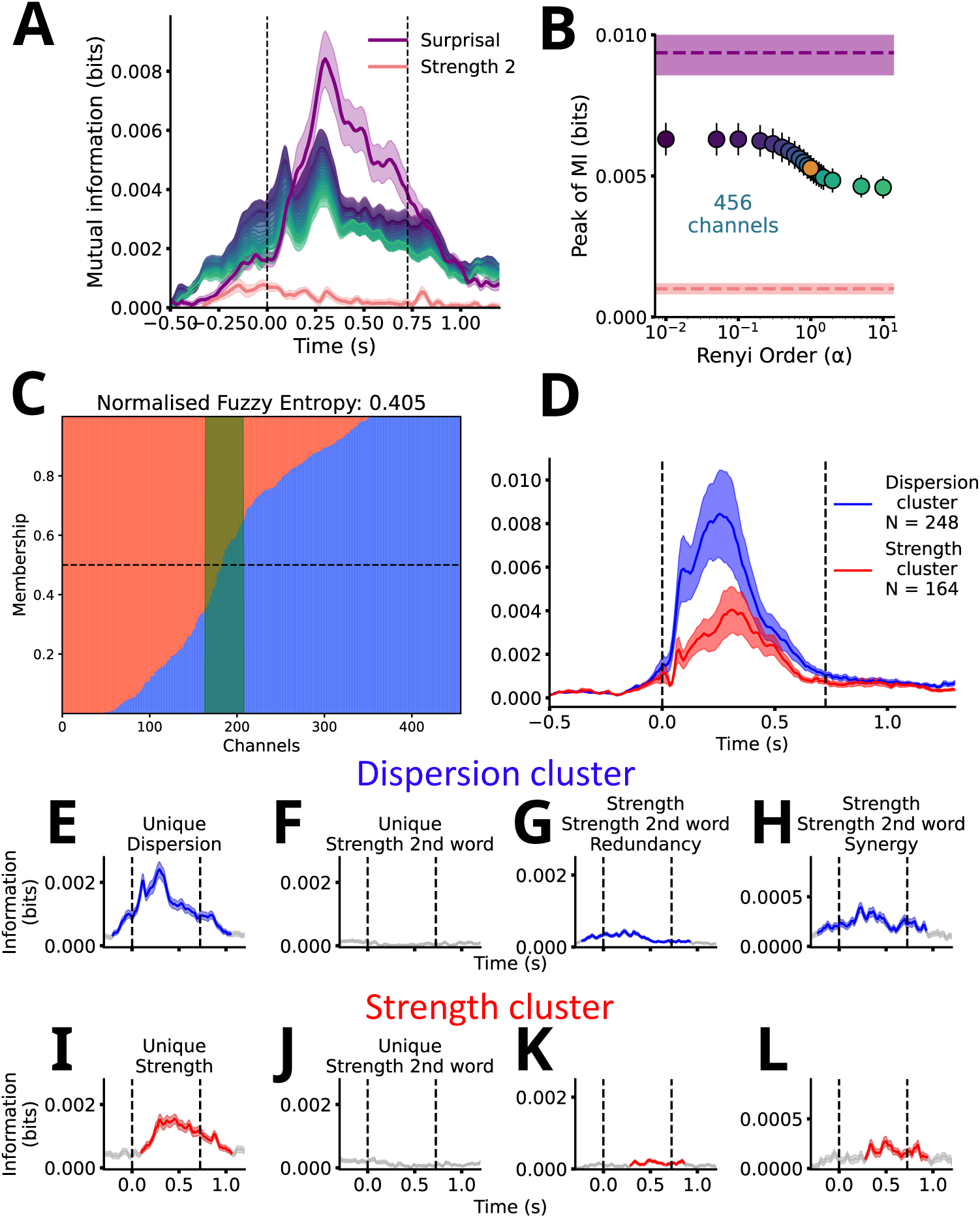
Extended Data 5. **(A,B)** Grand average of MI time courses across the 362 significant channels for different Rényi orders (*α*, color gradient), with a peak at 300 ms after word onset, on which the mutual information corresponding to surprisal(purple) and the entropy relative to the second most likely word(red), subbed Strength 2, were represented. Surprisal shows a clear peak around 300 ms while Strength 2 shows no activity. **(C)** Membership of the different channel relative to the dispersion(blue) and strength(red) clusters. In green, we represented the channels cut by the 0.65 threshold used in study. **(D)** Evoked response potential power relative to the onset of word for the dispersion(blue) and strength(red) clusters, averaged over all words. **E-L** PID measures computed between the strength associated to the second most likely word and either dispersion (**E-H**) or strength (**I-L**) on their respective clusters.

**Fig. 11.**
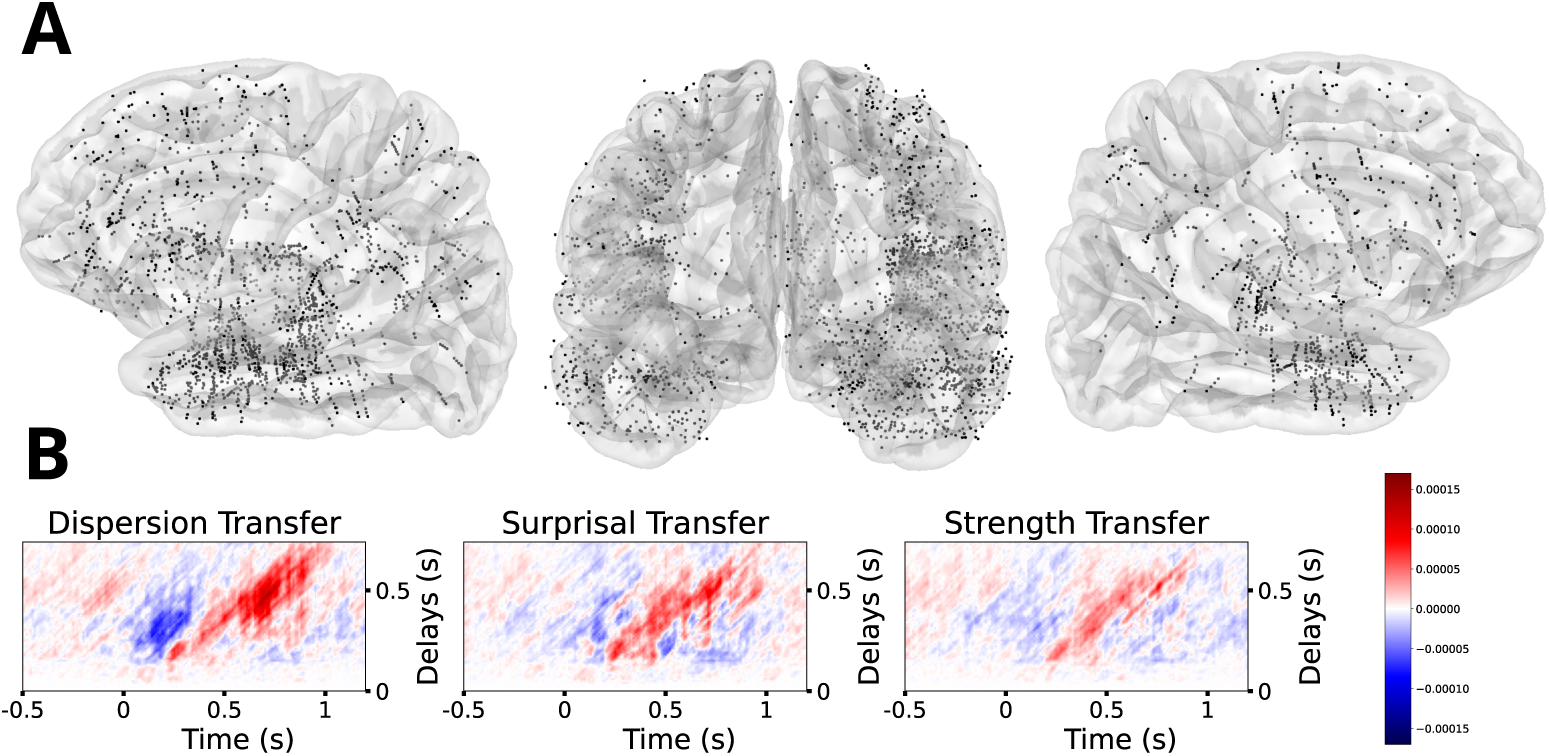
Extended Data 6. **(A)** Overall implantation of all subjects pooled together, resulting in 5194 channels. **(B)** Feature information transfer (FIT) time-delay matrices relative to dispersion, surprisal and strength.

**Fig. 12.**
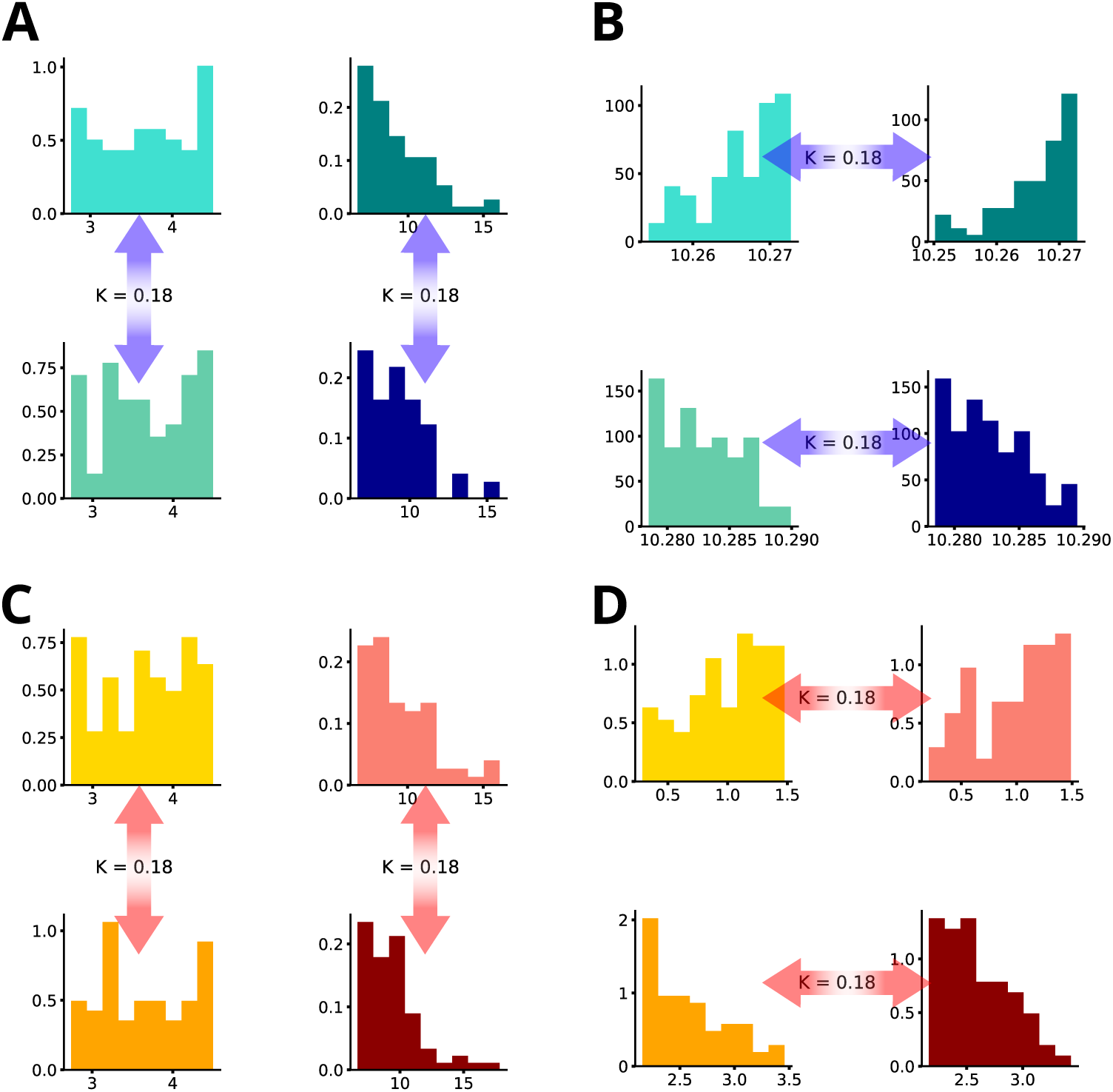
Extended Data 7. **(A)** Distribution of dispersion across the dispersion-surprisal quartiles. Two quartiles vertically aligned present similar dispersion values (K = 0.18). **(B)** Distribution of surprisal across the dispersion-surprisal quartiles. Two quartiles horizontally aligned present similar surprisal values (K = 0.18). **(C)** Distribution of strength across the strength-surprisal quartiles. Two quartiles vertically aligned present similar strength values (K = 0.18). **(D)** Distribution of surprisal across the strength-surprisal quartiles. Two quartiles horizontally aligned present similar surprisal values (K = 0.18).

**Fig. 13.**
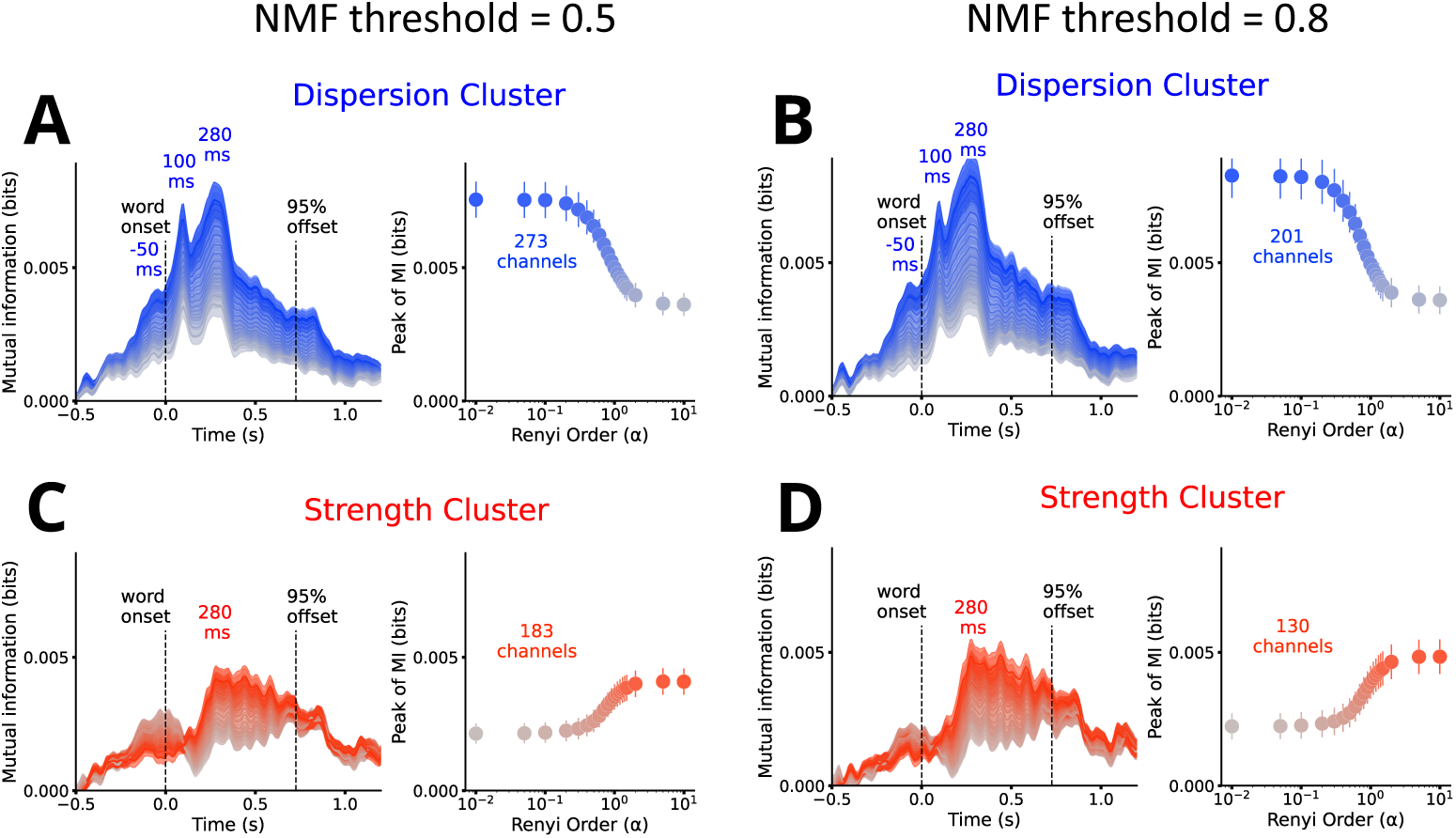
Extended Data 8. **(A-D)** Grand average of MI time courses across channels for different Rényi orders (*α*, color gradient) and MI peak as a function of Rényi order, averaged across channels. These were plotted for an NMF threshold of 0.5 for the dispersion **(A)** and strength **(C)** clusters, as well as for an NMF threshold of 0.8 for the dispersion **(B)** and strength **(D)** clusters.

**Fig. 14.**
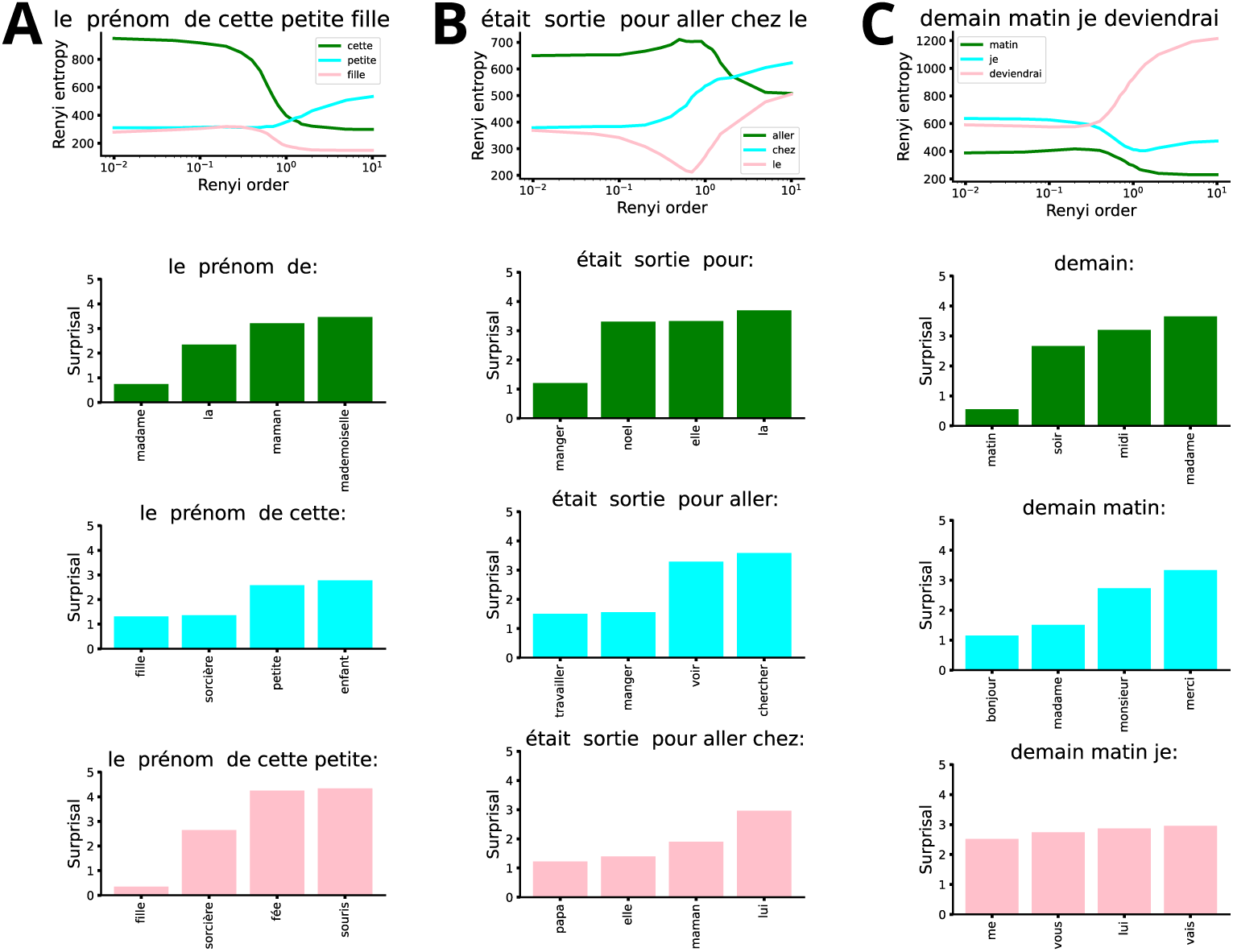
Extended Data 9. **(A-C)** Example of sentences extracted from the dataset and the rank of their associated Renyi entropy relative to the corpus for that same *α* parameter.

